# PGP1 personal genome assembly - a hybrid assembly dataset using ONT’s PromethION and PacBio’s HiFi sequencing

**DOI:** 10.1101/2021.09.03.458806

**Authors:** Hui-Su Kim, Changjae Kim, George Church, Jong Bhak

## Abstract

PGP1 is the first participant of Personal Genome Project. We present the PGP1’s chromosome-scale genome assembly. It was constructed using 255 Gb ultra-long PromethION reads and 97 Gb short paired-end reads. For reducing base calling errors, we corrected PromethION reads using 72 Gb PacBio HiFi reads. 327 Gb Hi-C chromosomal mapping data were utilized to maximize the assembly’s contiguity. PGP1’s contig assembly was 3.01 Gb in length comprising of 4,234 contigs with an N50 value of 33.8 Mb. After scaffolding with Hi-C data and extensive manual curation, we obtained a chromosome-scale assembly that represents 3,880 scaffolds with an N50 value of 142 Mb. From the Merqury assessment, PGP1 assembly achieved a high QV score of Q45.45. For a gene annotation, we predicted 106,789 genes with a liftover from the Gencode 38 and an assembly of transcriptome data.

## 1. Data description

We generated sequencing data of long-reads by ONT PromethION, short-reads by Illumina NovaSeq and Hi-C reads (Table 1). The sequencing data have been deposited in the NCBI database under BioProject: PRJNA734849. PGP1’s description is found also at http://genomics.org/PGP1. We collected PacBio HiFi reads from NCBI SRA accession SRX7671688. The *de novo* assembly of PGP1 genome is at chromosome-scale with a total length of 3.02 Gb, which consists of 3,880 scaffolds with 24 chromosomes and unplaced sequences (Table 2). Detailed features of the genome annotation are described in table 3. The sequence data and description are available at http://genomics.org/PGP1.

**Table 1.**
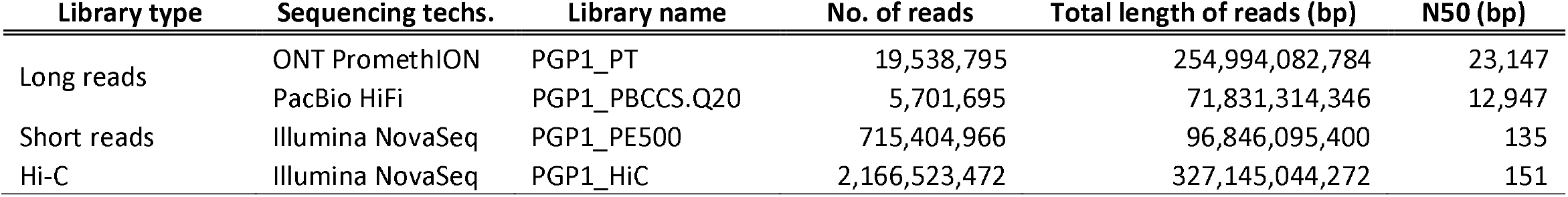
Statistics of long and short reads whole genome sequencing for PGP1.

| Library type | Sequencing techs. | Library name | No. of reads | Total length of reads (bp) | N50 (bp) |
| --- | --- | --- | --- | --- | --- |
| Long reads | ONT PromethION | PGP1_PT | 19,538,795 | 254,994,082,784 | 23,147 |
|  | PacBio HiFi | PGP1_PBCCS.Q20 | 5,701,695 | 71,831,314,346 | 12,947 |
| Short reads | Illumina NovaSeq | PGP1_PE500 | 715,404,966 | 96,846,095,400 | 135 |
| Hi-C | Illumina NovaSeq | PGP1_HiC | 2,166,523,472 | 327,145,044,272 | 151 |

**Table 2.** Statistics of PGP1 assembly.

|  | Contig assembly | Chromosome-scale assembly |
| --- | --- | --- |
| Contigs No. | 4,234 | 3,880 |
| Total length (bp) | 3,015,852,063 | 3,016,802,955 |
| N50 (bp) | 33,790,496 | 141,933,136 |
| Max contig length (bp) | 110,121,243 | 236,082,540 |
| Gap | 0.004% | 0.035% |
| GC contents | 40.87% | 40.87% |
| QV (from Merqury) |  | 45.4982 |
| Error rate (from Merqury) |  | 0.0000282 |

**Table 3.**
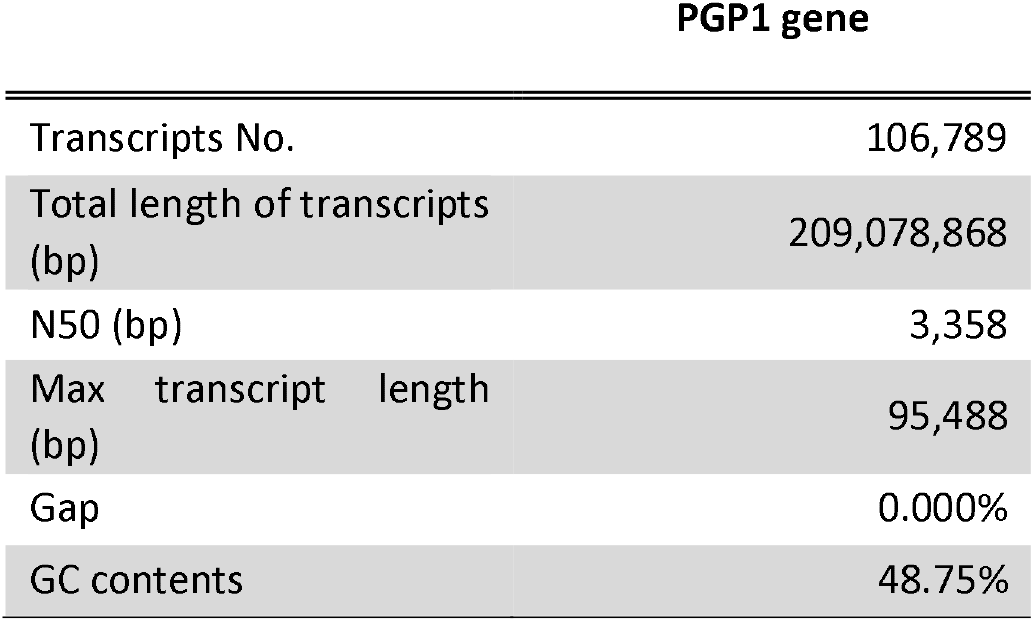
PGP1 genome annotation.

## 2. Experimental Design, Materials, and Methods

### 2.1. Sample preparation and whole-genome sequencing

DNA was extracted from samples from the PGP1 cell line from Coriell. For short-read sequencing, a 135 bp library was constructed, and the sequencing was conducted by Illumina’s NovaSeq platform. For long-read sequencing, we constructed libraries using the 1D ligation sequencing kit (SQK-LSK109), and the sequencing data was generated using ONT’s PromethION R9.4.1 platform. Base-calling was carried out using Guppy v3.5.4 with the Flip-Flop hac model. Libraries for Hi-C, the chromosome conformation capture data were generated using the Arima-Hi-C kit. The sequencing of Hi-C was performed using Illumina’s NovaSeq sequencer with a read length of 150 bp by Novogene.

### 2.2. Read preprocessing and whole genome assembly

Procedures are described in figure 1. Trimming adapter sequences and low-quality sequences in short reads were performed using Trimmomatic v0.36[1]. We used tadpole.sh program of BBtools suite v38.96 (https://sourceforge.net/projects/bbmap) for an error correction. Adapter sequences in PromethION reads were removed using Porechop v.0.2.4 (https://github.com/rrwick/Porechop), and we corrected PromethION reads against PacBio HiFi reads using Racon v1.4.3 program (https://github.com/isovic/racon).

**Fig. 1.**
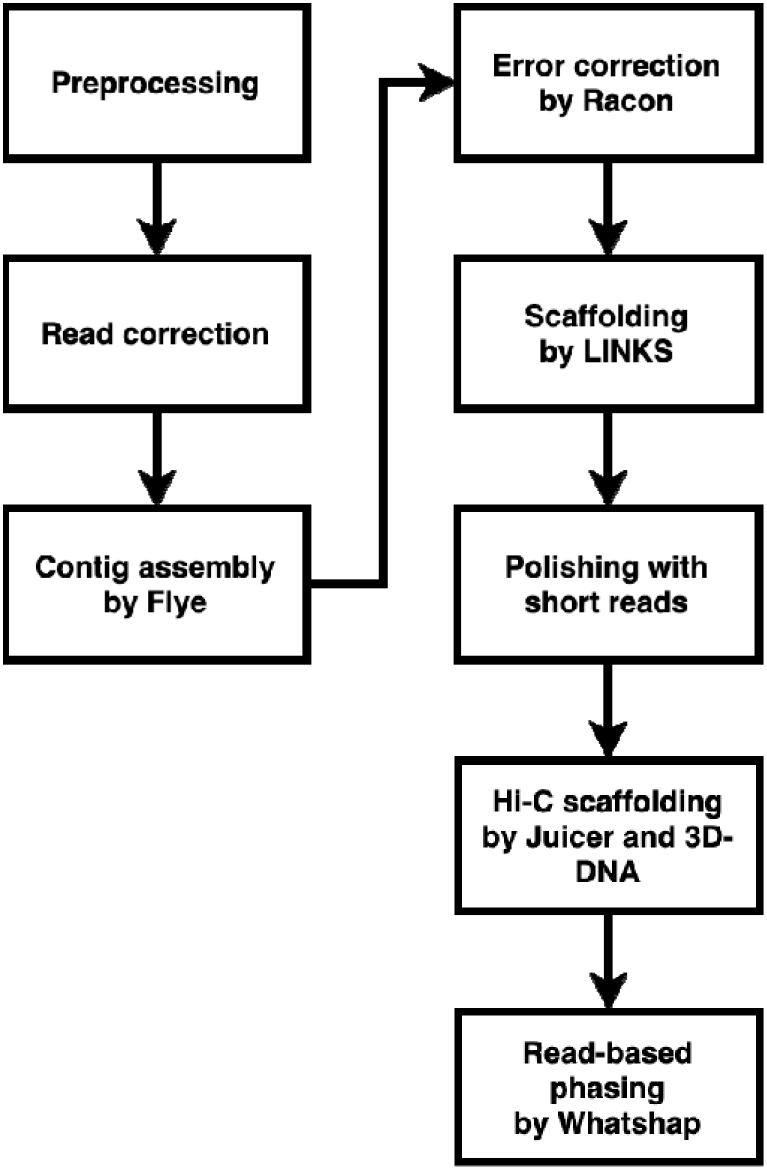
A bioinformatics pipeline for PGP1 genome assembly.

A *de novo* assembly was performed using Flye v2.5[2] program. Correcting base-errors in assembled contigs was conducted using Racon, and an extension of contigs using ultra-long PromethION reads was carried out using LINKS v1.8.7 (https://github.com/bcgsc/LINKS). Polishing the assembled contigs with short reads was performed using Pilon v1.23[3] twice.

For generating a chromosome-scale assembly, scaffolding contigs with Hi-C was performed using Juicer v1.5[4] and 3D-DNA pipeline[5]. For correcting mis-assemblies in the scaffolds, we used JBAT v1.11.08 program (https://github.com/aidenlab/Juicebox/wiki/Juicebox-Assembly-Tools) and corrected them manually. An assessment of PGP1 genome assembly was carried out using Merqury v1.3 program[6].

### 2.3. Read-based phasing and genome annotation

A read-based phasing of the assembly was performed using DeepVariant v1.1.0 (https://github.com/google/deepvariant) and WhatsHap v1.0[7], and we generated phased genome sequences from the phased variant-information using Bcftools v1.9 (http://github.com/samtools/bcftools). For gene annotation, a liftover of an annotated gene set from Gencode release 38 (https://www.gencodegenes.org/human/) using Liftoff v1.6.1[8] and a reference-guided transcriptome assembly using Stringtie v2.1.5 program[9] were conducted. The RNASeq data was obtained from SRA no. SRX683721, SRX683722, SRX683723.

## Specifications Table

|  |  |
| --- | --- |
| **Subject** | Biology |
| **Specific subject area** | Genomics |
| **Type of data** | Sequencing raw reads, Assembly, Tables, Figure |
| **How data were acquired** | PromethION flow-cell R9.4.1 (Oxford Nanopore Technologies)<br>PacBio HiFi (Pacific Bioscience)<br>NovaSeq (Illumina)<br>Hi-C (Arima-Genomics) |
| **Data format** | Raw reads (fastq), Assembly (fasta), Protein and Transcript sequences (fasta), Genome annotation (gff3) |
| **Parameters for data collection** | DNA from the PGP1 cell line used for library preparation and sequencing. |
| **Description of data collection** | Total genomics DNA extraction was performed using DNeasy Blood & Tissue Kit from QIAGEN. The library construction and whole genome sequencing were performed using Illumina's NoveSeq (100bp x2, short reads), ONT's PromethION (long reads), and Illumina's NoveSeq platform (151bp x2, Hi-C). |
| **Data source location** | Institution: Korean Genomics Center (KOGIC)<br>City/Town/Region: Ulsan city<br>Country: Korea, republic of |
| **Data accessibility** | Raw data was deposited in the NCBI database under BioProject: PRJNA734849 (https://www.ncbi.nlm.nih.gov/bioproject/PRJNA734849), SRA: SRX11055733, SRX11055734, SRX11055735, SRX11055736, SRX11055737, SRX11055738 (https://www.ncbi.nlm.nih.gov/sra?linkname=bioproject_sra_all&from_uid=734849).<br>Its description, the final assembly and data information are also found at http://genomics.org/PGP1. |

## Value of the Data

- This is a *de novo* genome assembly of PGP1 of Personal Genome Project of Harvard Medical school.
- The genome assembly of a male Caucasian and sequencing data add to current genomic representation of North-Eastern Europeans and is a useful resource for further in-depth analyses of European genomic structure and diversity in higher resolution.
- We share a hybrid assembly pipeline used in this study for constructing a high-quality chromosome-scale assembly from PromethION, PacBio-HiFi, and Hi-C data which can be a useful approach for the bioinformatics community specializing in genome assembly.

## Declarations

### Ethics Statement

This study was a part of Korean Personal Genome Project (KPGP also known as PGP-Korea) and was approved by the Institutional Review Board at Genome Research Foundation with IRB- REC- 20101202 - 001. The anonymous sample donor has signed a written informed consent to participate in the whole genome sequencing and following analysis in compliance with the Declaration of Helsinki.

### Consent for publication

The (KPGP) informed consent included a section about data publication, which was consented to.

### CRediT author statement

**Hui-Su Kim:** Methodology, Formal analysis, Writing - Original Draft, Data Curation, Visualization, and Editing. **Jong Bhak:** Funding acquisition, Project administration, Supervision, Resources, Conceptualization, and Writing - Review & Editing. **Changjae Kim**: Performed DNA preparation, sequencing, and data quality checking. **George Church**: Initiated and supervised PGP, provided cell-line. All authors read and approved the finalized manuscript.

## Acknowledgements

We thank GenomeLab, PGI of GRF, and KOGIC members for providing technical assistance and discussions. We also thank the Korea Institute of Science and Technology Information (KISTI) that provided us with the Korea Research Environment Open NETwork (KREONET).

